# Transposon-induced inversions activate gene expression in Maize pericarp

**DOI:** 10.1101/2021.03.08.434479

**Authors:** Sharu Paul Sharma, Tao Zuo, Thomas Peterson

## Abstract

Chromosomal inversions can have considerable biological and agronomic impacts including disrupted gene function, change in gene expression and inhibited recombination. Here we describe the molecular structure and functional impact of six inversions caused by Alternative Transpositions between *p1* and *p2* genes responsible for floral pigmentation in maize. In maize line *p1-wwB54*, the *p2* gene is expressed in anther and silk but not in pericarp, making the kernels white. We identified inversions in this region caused by transposition of *Ac* and *fractured Ac* (*fAc*) transposable elements. These inversions change the position of a *p1* enhancer and activate the expression of *p2* in the kernel pericarp, resulting in red kernel color. We hypothesize that these inversions place the *p2* gene promoter near a *p1* gene enhancer, thereby activating *p2* expression in kernel pericarp.

## Introduction

Transposable elements are segments of DNA that can move within a genome. The maize *Activator* (*Ac*) and *Dissociation* (*Ds*) transposable elements are members of the *hAT* transposon super-family, which is widespread in eukaryotes (Rubin *et al*. 2001). Barbara McClintock discovered these transposons while studying the phenomenon of chromosome breakage. She identified *Ds* as a locus on the short arm of chromosome 9 in some maize stocks where chromosome breaks occurred frequently. She also showed that *Ds* is dependent on another element *Ac* which is autonomous and can itself transpose. The *Ac/Ds* system was also reported to induce a variety of chromosomal rearrangements, such as translocations, deletions, duplications, and inversions (McClintock 1950, 1951). The autonomous *Ac* element is 4565 bp in length and carries a complete transposase gene. *Ds* elements vary in size and internal sequence and lack a functional transposase gene, making them non-autonomous (Lazarow *et al*. 2013). The *Ac* transposase is known to bind to subterminal motif sequences of *Ac/Ds* elements and then cut at the transposon 5’ and 3’ TIRs (Terminal Inverted Repeats; 11 bp imperfect repeats) (Becker and Kunze 1997). *Ac* transposase can recognize and act on the termini of a single element (Standard Transposition), or the termini of two different elements (Alternative Transposition); for example, the 5’ end of *Ac* and the 3’ end of a second nearby element such as *Ds* or *fractured Ac* (*fAc*) (Su *et al*. 2018). Standard Transposition events change only the position of a single element, while Alternative Transposition events can produce a variety of genome rearrangements, depending on the relative orientations of the TE termini and the location of the target site. When two transposons are in direct orientation, the internal-facing termini are present in a reversed orientation compared to the termini of a single transposon. In this configuration, the two facing termini can undergo Reversed-Ends Transposition (RET) (Huang and Dooner 2008; Zhang and Peterson 2004; Zhang *et al*. 2009) to induce deletions (Zhang, J. and Peterson 2005; Zhang, J. *et al*. 2006), duplications (Zhang *et al*. 2013), Composite Insertions (Zhang *et al*. 2014; Su *et al*. 2018, 2020), inversions (Zhang and Peterson 2004; Yu *et al*. 2011) and reciprocal translocations (Pulletikurti *et al*. 2009; Zhang *et al*. 2009). For example, Zhang *et al*. 2009 described 17 reciprocal translocations and two large pericentric inversions derived by RET from a progenitor allele containing *Ac* and *fAc* insertions in the maize *p1* gene. The frequent occurrence of these structural changes and the fact that *Ac* inserts preferentially in or near genic regions (Kolkman *et al*. 2005) suggest that Alternative Transposition events may have a significant impact on the genome and transcriptome. Additionally, inversions provide an opportunity to analyze the function of *cis*-regulatory elements, such as enhancers, in a native (non-transgenic) context.

The maize *p1* and *p2* genes are closely linked paralogous genes located on the short arm of chromosome 1 that originated by duplication of an ancestral *P*^*pre*^ gene, approximately 2.75 mya (Zhang, P. *et al*. 2000). These genes are separated by a ∼70 kb intergenic region and coincide with a major QTL for levels of silk maysin, a flavone glycoside with antibiotic activity toward corn earworm (Zhang *et al*. 2003; Meyer *et al*. 2007). Both *p1* and *p2* encode highly similar R2R3 Myb transcription factors involved in controlling the structural genes *c2, chi*, and *a1*, encoding chalcone synthase, chalcone isomerase, and dihydro-flavonol reductase, respectively (Dooner *et al*. 1991; Grotewold *et al*. 1994). These enzymes of the flavonoid biosynthetic pathway produce red phlobaphene pigments in maize floral organs. *p1* is expressed in maize kernel pericarp, cob, and silk, while *p2* is active in anther and silk (Zhang, P. *et al*. 2000; Goettel and Messing 2009). Different *p1* alleles are indicated by a two-letter suffix indicating their expression in kernel pericarp and cob glumes; for example, *p1-ww* specifies white (colorless) pericarp and white cob, while *P1-wr* indicates white pericarp and red cob.

The robust visual phenotypes and abundance of alleles with *Ac* insertions (Athma *et al*. 1992; Moreno *et al*. 1992) make the *p1/p2* cluster an ideal genetic system to analyze the genetic impact of Alternative Transposition events. The *p1-wwB54* allele has a deletion of *p1* exons 1 and 2 along with insertions of *Ac* and fractured *Ac* (*fAc*) elements upstream of *p1* exon 3 (Yu *et al*. 2011). Because exons 1 and 2 encode most of the essential Myb DNA binding domain (Grotewold *et al*. 1991) their deletion renders the *p1* gene non-functional leading to white kernel pericarp and white cob. The 5’ *Ac* and 3’ *fAc* termini are in a reversed orientation, separated by a 331 bp inter-transposon segment. These elements exhibit frequent RET, leading to chromosome breakage and rearrangements such as deletions and inversions (Yu *et al*., 2011). Here, we used the *p1-wwB54* allele as a starting point to isolate a variety of *p1/p2* gain of function alleles. Among these, we identified independent cases of inversions with varying degrees of red kernel pigmentation, possibly due to the activation of *p2* in pericarp tissue. Here we describe the detailed structures and *p2* expression characteristics of six inversion cases.

## Materials and Methods

### Screening for Inversions derived from RET

The inversion alleles described here were derived from *p1-wwB54* (Figure 1). Stock J (*p1-ww*[*4Co63*] *r1-m3::Ds*) (described in Zhang *et al*. 2003) was used as common genetic background and to detect the presence of *Ac* by excision of *Ds* from *r1-m3*. The occurrence of red kernel pericarp in *p1-wwB54* was used as a visual screen for *p2* activation in the pericarp (see Materials and Methods in Su *et al*. 2020). *p1-wwB54* has white kernels, but approximately 1 in 8 ears were found to have a single red kernel, and ∼1 in 40 ears had a multi-kernel red sector (Figure 1, F86). The occurrence of a sector of red-colored pericarp on single or multiple kernels reflects the stage of ear and kernel development at which an activating mutation (e.g. transposition) occurred. Events that occurred sufficiently early (prior to embryo formation) can be inherited (Emerson 1917). The red kernels were selected and planted, and in cases where the new structure was transmitted through meiosis, the resulting plants would produce whole ears with red kernels (Figure 1, S25). The pericarp is maternal tissue and hence the red color phenotype is independent of the pollination parent.

**Figure 1:**
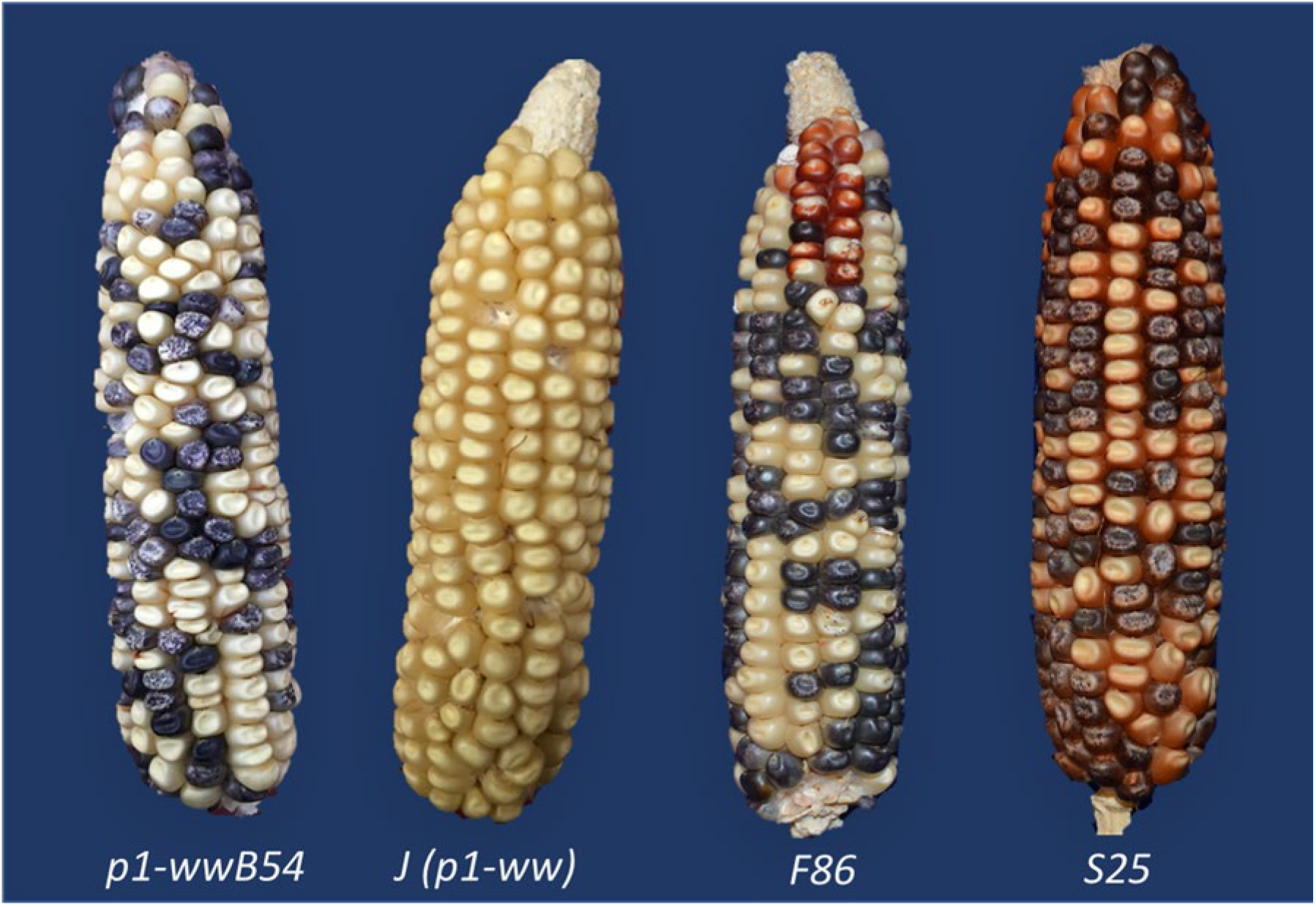
Ears of Different Maize *p1/p2* Alleles. Alleles *p1-wwB54* and *J (p1-ww)* have white (colorless) kernel pericarp. F86 is a *p1-wwB54* ear in which a sector of kernels near the ear tip has red pericarp due to activation of *p2*. S25 is an inversion allele with red pericarp color on the whole ear. Kernels with purple-sectored aleurone are due to *Ac*-induced excision of *Ds* from *r1-m3::Ds*.

### DNA extraction, Gel Electrophoresis, and Southern Blotting

Genomic DNA was extracted from maize seedling leaves by a modified CTAB method (Saghai-Maroof *et al*. 1984) and digested with different restriction enzymes according to the manufacturer’s instructions. For Southern blotting, genomic DNA digests were done with *KpnI, HpaI*, and *EcoRV*. Agarose gels (0.7%) were run under 30 to 50 volts for 18 to 24 hours for maximum separation of large fragments. The DNA was then transferred to a membrane for 24 hours, followed by probing each membrane with Fragment-15 (*f15*), a 411 bp sequence two copies of which are located within the enhancer of the *p1* gene (Zhang F. and Peterson 2005).

### PCR, iPCR, and Sequencing

PCR was performed with 20 µL reaction volumes under the following temperature conditions: 95° for 2 min, then 35 cycles at 95° for 30 sec, 60° for 30 sec, and 72° for 1 min per 1-kb length of the expected PCR product, then final extension at 72° for 5 min. For initial PCR screening of new alleles, a high-efficiency agarose gel electrophoresis method was used to visualize PCR products (Sharma and Peterson 2020). Inversion breakpoint junctions ending with *fAc* elements were obtained by inverse-PCR (iPCR; Ochman *et al*. 1988). Inversion breakpoints at *Ac* elements were isolated by *Ac* casting (Singh *et al*. 2003; Wang and Peterson 2013). This method relies on the occurrence of frequent *Ac* transpositions to closely linked sites during plant development. For each inversion, genomic DNA was isolated from seedling leaf tissue and then the region containing the breakpoint was amplified by two pairs of nested PCR primers (Set 1 and then Nested in Table S1). The inversion breakpoint regions from I-PCR and *Ac* casting were sequenced by the Iowa State University DNA Sequencing Facility. Sequences were analyzed using Snapgene (snapgene.com) and BLAST (Zhang, Z. *et al*. 2000).

### RT-PCR Detection of p2 Expression

Pericarps were peeled from kernels 15 to 20 days after pollination (DAP) and flash-frozen in liquid nitrogen. Three biological replicates (pericarps from 3 sibling ears) were pooled to extract RNA. RNA was isolated using Purelink Plant RNA Reagent, treated with NEB DNaseI, and reverse transcribed to cDNA using Invitrogen^™^ SuperScript^™^ II Reverse Transcriptase kit using protocols recommended by the product suppliers. Two technical replicates of reverse transcription were used per sample. cDNAs were amplified by PCR using primers specific to exons 1 and 3 of the *p2* gene transcript (Table S3). Primers specific to the maize *Beta-tubulin* gene were used as an internal control.

## Data availability

Maize genetic stocks are available by request to T.P. Sequences reported here are available in the Supplemental Material.

## Results

Due to the deletion of *p1* exons 1 and 2, the *p1-wwB54* was expected to be a stable null allele. We were surprised to see ears carrying *p1-wwB54* produced red kernel pericarp sectors of varying sizes (Figure 1). We hypothesized that the *p2* gene, which is normally not expressed in kernel pericarp, could be activated by inversions generated by Reversed Ends Transposition (RET) (Zhang and Peterson 2004; Zhang, J. and Peterson 2005; Zhang *et al*. 2009, 2013; Huang and Dooner 2008; Yu *et al*. 2011; Su *et al*. 2020). A diagram of this model showing an inversion with breakpoints in the *p2* promoter region is shown in Figure 2. According to this model, RET would begin with excision of the *Ac* 5’ end and *fAc* 3’ end in *p1-wwB54*, followed by insertion of the excised termini into a new target site unique for each event (Figure 2, *a/b*). If the 5’ end of *Ac* (solid red arrowhead, Figure 2) joined with the ‘*a*’ side of the target sequence, and 3’ end of *fAc* (white arrowhead, Figure 2) joined with the ‘*b*’ side of the target site, the segment from 5’ end of *Ac* up to the target site *a/b* will be inverted (for animation, see Supplemental data). The resulting structure (Figure 2, *Lower*) contains an inversion of the *p1-p2* interval; if the *p2* gene promoter region is inserted sufficiently near the *p1* 3’ pericarp enhancer (Sidorenko *et al*., 2000), *p2* may be expressed in the kernel pericarp.

**Figure 2:**
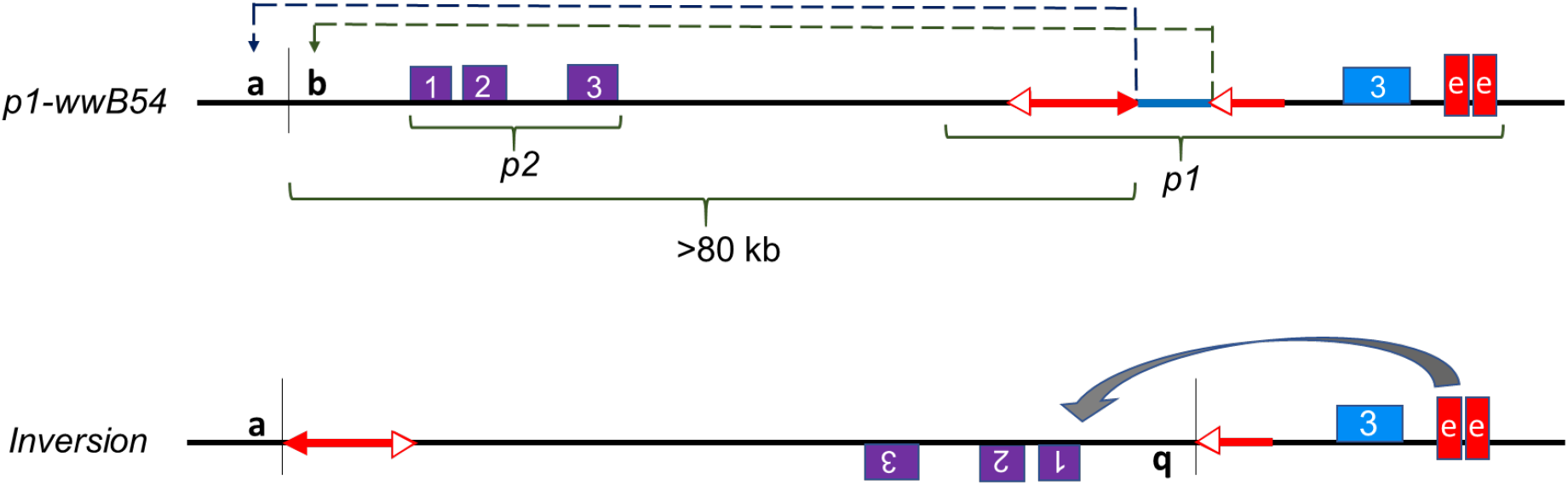
Model of RET-induced inversion leading to *p2* activation. *Upper: Diagram of progenitor allele p1-wwB54 and nearby p2 gene:* Purple and blue boxes indicate exons of *p2* and *p1* genes, respectively. Red arrows represent *Ac* (with two arrowheads) and *fAc* (with single arrowhead) elements. Red boxes indicate two copies of *p1* enhancer fragment *f15*. Dashed lines indicate *Ac/fAc* excision by RET and re-insertion at *a/b* target site upstream of *p2*. The 331 bp DNA fragment between *Ac* and *fAc* (blue line) is lost during the transposition event. The same symbols and coloring scheme are used in other figures in this paper. *Lower: Inversion:* Inverted segment extends from point *a* (*Ac* junction) to point *b* (*fAc* junction) and includes *Ac, p1-p2* intergenic region, and *p2* gene. In the inversion allele, the proximity of the *p2* promoter to the *p1* 3’ enhancer may activate *p2* expression in the pericarp.

### Screening for Inversions

To obtain RET-induced inversions, ears from several thousand plants carrying the *p1-wwB54* allele were screened for kernels with red pericarp (example in Figure 1, third ear from left). Selected red kernels were grown and propagated to obtain stable lines with various shades of red kernel pericarp. Genomic DNA preparations from these lines were tested for structural rearrangements by PCR using sets of primer pairs (Table S1) that can amplify the *Ac* and *fAc* junctions in *p1-wwB54:* primer set 1 detects the *p1/3’ Ac* junction, and primer set 2 detects the *5’ Ac/p1/3’ fAc* segment (Figure 3A). Simple *Ac* transposition or RET-induced deletion (Yu *et al*., 2011) would result in negative PCR for both sets 1 and 2; while the formation of Composite Insertions (Su et al., 2020) results in retention of both junctions. Whereas, RET-induced inversion would result in retention of the *p1/3’ Ac* junction (Set 1 positive), and loss of the *5’ Ac/p1/3’ fAc* segment (Set 2 negative). Using this test, several cases of putative inversions were detected (Figure 3B). These cases were further tested using primers flanking the downstream *fAc/p1* junction (Table S1) to confirm the retention of *fAc* at its original position next to *p1* exon 3. Following confirmation of potential inversions, the new *Ac* and *fAc* inversion breakpoint junction sequences (*a/Ac* and *fAc/b* in Figure 3A) were amplified from genomic DNA using direct PCR, *Ac* Casting, or iPCR (see Methods) along with nested PCR. Once obtained, both inversion breakpoint junctions were sequenced (list of primers in Table S2). Junction sequences were examined to confirm expected orientations based on the established *p1* and *p2* genomic sequence data (Zhang *et al*. 2006) and the presence of 8 bp TSDs (Target Site Duplication) characteristic of *Ac* transposition (Figure 3A, yellow box and Table S4).

**Figure 3:**
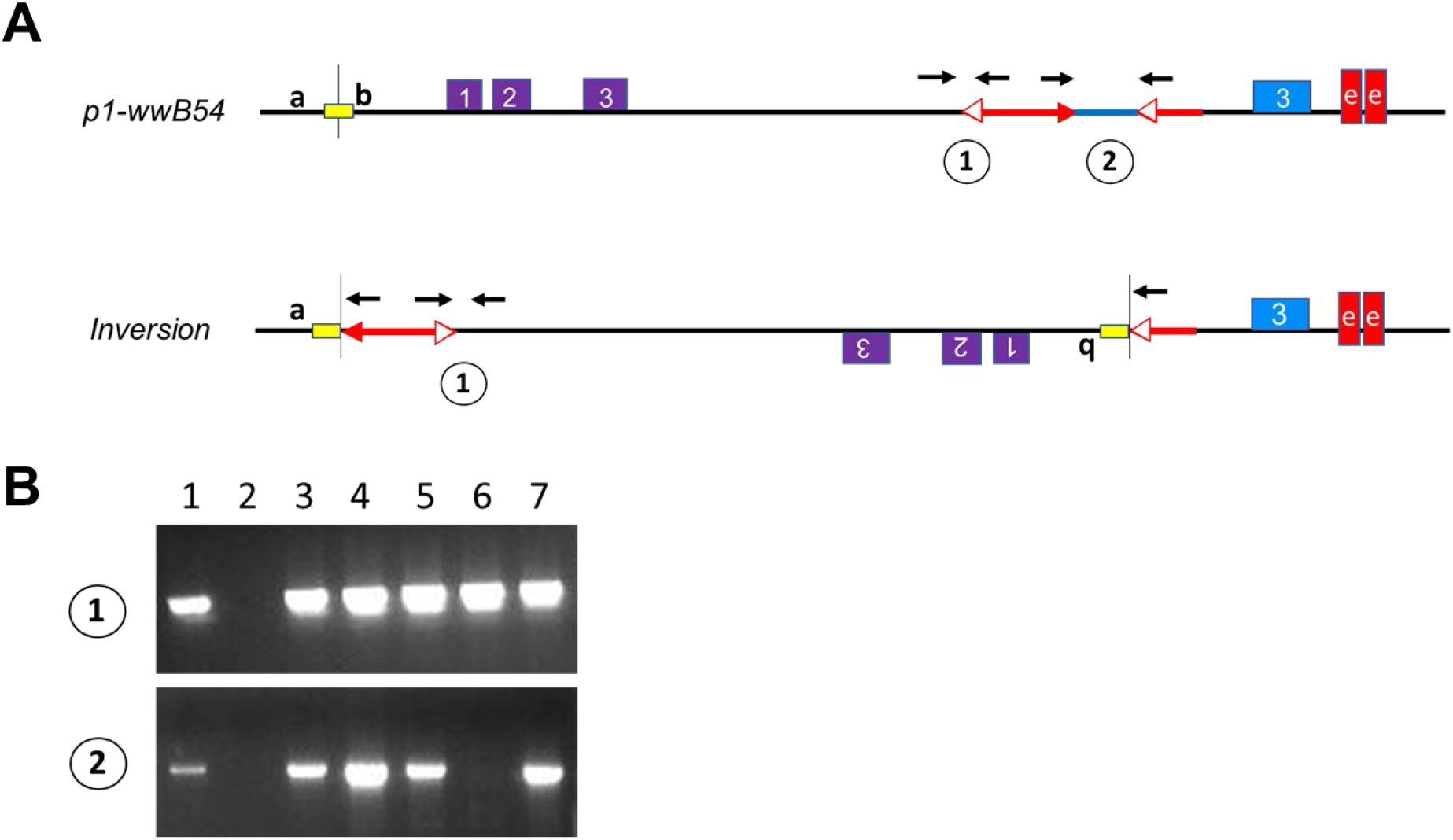
PCR Test for Inversions. A) Progenitor *p1-wwB54* and derived *Inversion* allele structures showing locations of primers (black arrows) used in PCR tests. Primer Set 1 detects the *p1-Ac* junction which is present in both *p1-wwB54* and *Inversion;* Primer Set 2 detects the *Ac/p1/fAc* segment which is present in *p1-wwB54* and absent in *Inversion*. Yellow box is the 8 bp target site duplicated in inversion. B) Agarose Gel image showing an example PCR using Primer Set 1 (upper) and Primer Set 2 (lower). Lane 1, positive control (*p1-wwB54*); Lane 2, negative control (*p1-ww* Stock J); Lanes 3 – 7, candidates tested. Only lane 6 (allele *132, not one of the cases described here*) is positive for Set 1, negative for Set 2, as expected for inversions.

### Structure of Inversions

The structures of six independent inversions with red kernel pericarp were determined. Ears produced by plants carrying these inversions are shown in Figures 1 and 4. The inversion junctions were PCR-amplified and sequenced as described above, and their sequences compared with established *p1* and *p2* genomic sequences to identify the breakpoint locations. One breakpoint common to all cases is at the 5’ end of *Ac* (Figure 5, vertical blue line), as expected for inversions originating by RET of *Ac* and *fAc* elements. The second breakpoint unique to each allele is at the transposition target site, located in a ∼1 kb window from 2.6 to 3.5 kb upstream of the *p2* transcription start site in these six cases (Figure 5, vertical red lines). These inversions reduce the distance between the *p2* transcription start site and the *p1* enhancer from 83.3 kb in the parental *p1-wwB54* allele to less than 10 kb in the inversion alleles (Figure 5 and Table S4). The inverted fragment size ranges from 80.9 to 81.8 kb. Each inversion allele contains an 8 bp repeat sequence at the inversion junctions, precisely at the ends of the *Ac* and *fAc* termini (Table S4). These 8 bp repeats represent the signature Target Site Duplications (TSDs) resulting from the staggered DNA cut made by *Ac* transposase. The presence of matching breakpoint TSDs confirms that each inversion originated from a single Alternative Transposition event involving the *Ac/fAc* elements.

**Figure 4:**
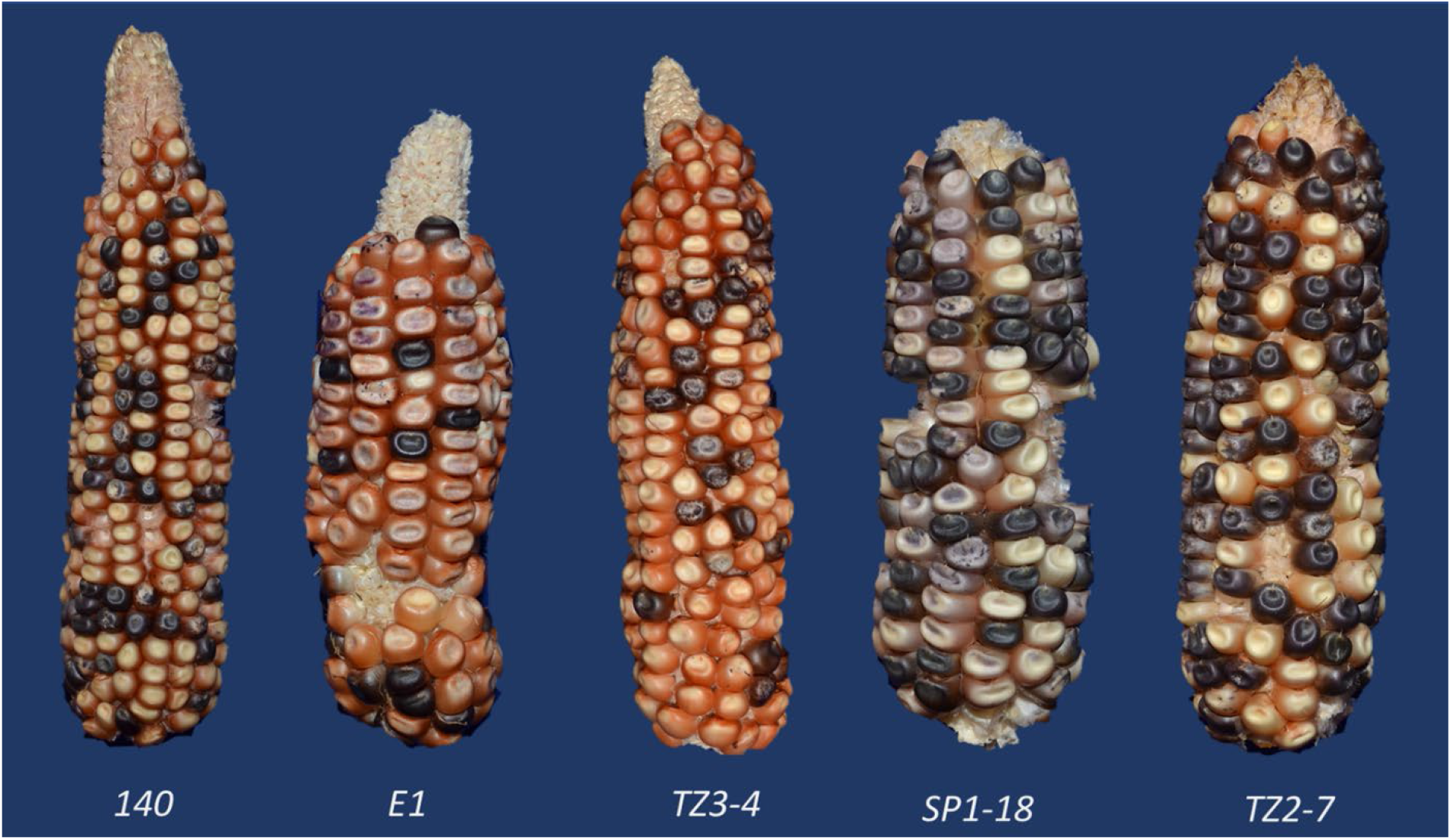
Representative Ears of Five Inversion Alleles. Ears have varying shades of red kernel pericarp due to *p2* activation. The sixth inversion case S25 is shown in Figure 1. Some kernels have purple or purple-sectored aleurone due to *Ac*-induced excision of Ds from *r1-m3* leading to anthocyanin pigmentation.

**Figure 5:**
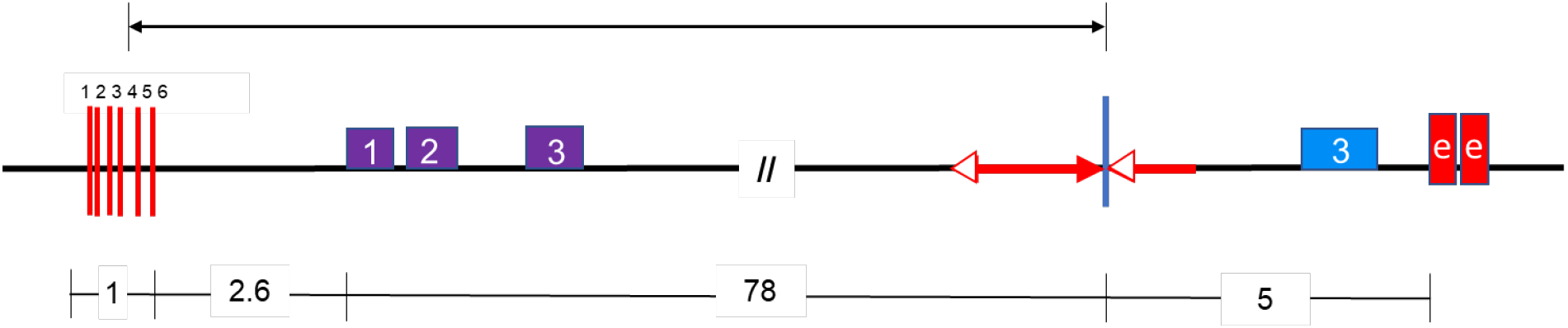
Map of the six inversion alleles. The vertical blue line is one breakpoint, and the red lines indicate the second breakpoint unique to each inversion. Numbers on red lines correspond to alleles, 1, 140; 2, E1; 3, TZ3-4; 4, SP1-18; 5, S25; 6, TZ2-7. Numbers below the figure are distances.

After identifying the endpoints of the inversions, Southern blotting experiments were conducted to examine the internal structures of the inverted fragments. Endonuclease *Kpn*I has recognition sites located such that the unique inversion breakpoint and the *p1* enhancer are contained in the same restriction fragment in all six cases of inversions (Figure 6). This inversion junction fragment was detected by hybridization with fragment-15 (*f15*) from within the *p1* enhancer. As shown in Figure 6A, *P1-rr4B2* (lane 3) gives two bands of size 6.3 kb and 8.6 kb as expected because it has two copies of the enhancer, one on each side (5’ and 3’) of the *p1* gene (Figure 6B; Sidorenko *et al*., 2000). Whereas the inversion progenitor *p1-wwB54* (lane 4) gives a single band of 13.5 kb representing the 3’ enhancer fragment; the 5’ enhancer fragment is deleted in this allele (Figure 6C). The six inversion alleles (lanes 5 – 10) have progressively decreasing band sizes, ranging from 12 to 10.5 kb, reflecting the size differences resulting from different junction breakpoints ‘*b*’ in each inversion (Figure 6D). Similar results were obtained using other restriction enzymes including *Hpa*I and *Eco*RV (Figure S1) and probes (*Ac-H* for the *Ac* element, and *p1* Fragment *8b* for *p2* intron 2; not shown). All the results are consistent with the presence of a simple inversion in each of these six cases, with no evidence of additional rearrangements.

**Figure 6:**
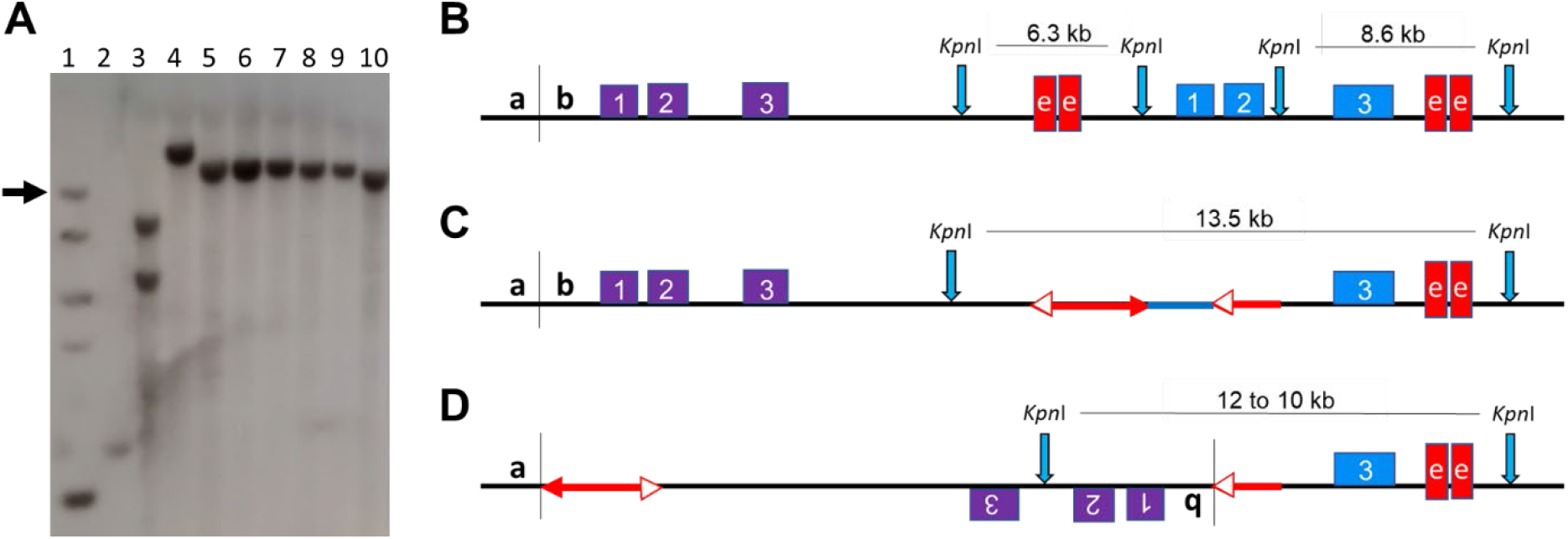
Genomic Southern Blot Analysis of Inversion Alleles. A) Southern blot of genomic DNA samples from inversion homozygotes digested with *Kpn*I and probed with fragment *f15* from the *p1* enhancer (red boxes in B, C and D). Lane 1, DNA ladder (arrow points to 10 kb band); Lane 2, J (*p1-ww*); Lane 3, *P1-rr4B2*; Lane 4, progenitor *p1-wwB54* (top band is 13.5 kb); Lane 5, 140; Lane 6, E1; Lane 7, TZ3-4; Lane 8, SP1-18; Lane 9, S25; Lane 10, TZ2-7. The six inversions (Lanes 5 – 10) are arranged in order of decreasing band sizes (from 12 to 10.5 kb). B, C and D) Diagrams showing *Kpn*I restriction sites (vertical blue arrows) in B) *P1-rr4B2*, C) progenitor *p1-wwB54* and D) inversions. Southern blot band sizes reflect differences in the sites of *fAc* insertion in the *p2* promoter (breakpoint *b*).

## p2 Expression in Inversions

The expression of the *p2* gene in plants homozygous for the inversion alleles was analyzed by RT-PCR. RNA was extracted from pericarps of homozygous plants collected 15-20 DAP (days after pollination) (Figure 7). *P1-rr4B2* was used as a positive control (Figure 7, lane 2) because *p1* is expressed in *P1-rr4B2* pericarp and the same *p2* primers can amplify *p1* transcripts due to sequence similarity. The six inversion alleles were derived from the *p1-wwB54* maize line which has a deleted *p1* gene and intact *p2* gene. The *p2* gene transcript was not detected in the pericarp tissue of *p1-wwB54* (Figure 7, lane 3), confirming previous results that *p2* is normally not expressed in kernel pericarp (Zhang, P. *et al*. 2000). However, *p2* transcripts were seen in all six inversion cases (Figure 7, lanes 4 – 9). To confirm the origin of these transcripts, the RT-PCR product of one inversion was sequenced and found to have sequence polymorphisms matching the *p2* gene (Figure S2). These results show that, unlike the progenitor *p1-wwB54, p2* is expressed in the pericarp tissue of all six inversion alleles. This ectopic *p2* expression likely resulted from the proximity of the *p2* gene promoter within the inverted fragment to the *p1* 3’ enhancer. In the progenitor *p1-wwB54*, the *p2* promoter region and *p1* 3’ enhancer are separated by 83.3 kb, whereas in the inversion alleles, this distance was reduced to between 7.4 and 8.2 kb. These results demonstrate the unique ability of inversions to modify gene expression near inversion breakpoints by changing the distance from regulatory elements to their target genes.

**Figure 7:**
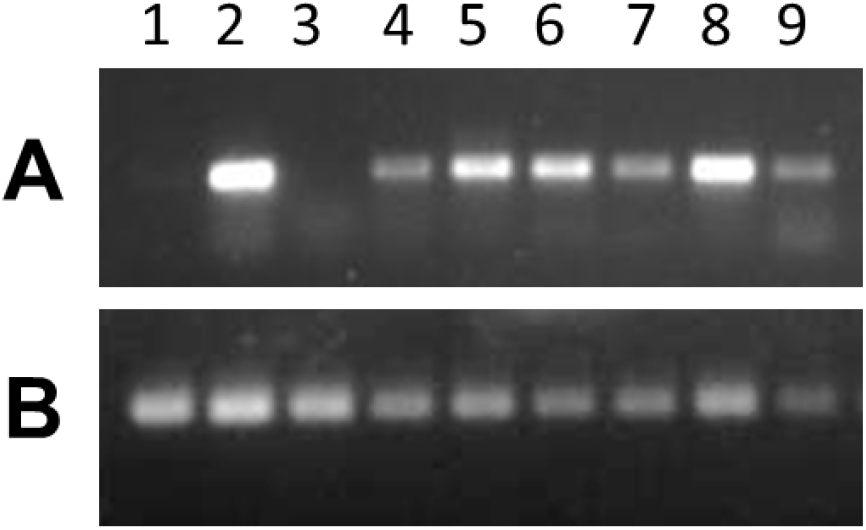
RT-PCR. Agarose gel images showing RT-PCR results using RNA extracted from pericarp tissue and reverse transcribed to cDNA. PCR with primers from A) *p2* exons 1 and 3, B) *Beta-tubulin* as an internal control. Lane 1, J (*p1-ww*) is negative control; Lane 2, *P1-rr4B2* is positive control for *p1* expression; Lane 3, *p1-wwB54* is the progenitor and lacks *p2* expression; Lane 4, 140; Lane 5, E1; Lane 6, TZ3-4; Lane 7, SP1-18; Lane 8, S25; and Lane 9, TZ2-7. All six inversion alleles are positive for *p2* expression.

## Discussion

### Mechanisms of inversions

A variety of molecular mechanisms are known to induce inversions. The double-strand break (DSB) mechanism involves breakage and then repair by Non-Homologous End Joining (NHEJ) (Moore and Haber 1996). If two double-strand breaks occur on the same chromosome, re-ligation of the DNA molecule via NHEJ can form inversions (Hefferin and Tomkinson 2005), deletions, or inversions flanked by inverted duplications, if the DSBs are staggered cuts (Ranz *et al*. 2007). Additionally, inversions can result from ectopic recombination (Non-Allelic Homologous Recombination, NAHR) between dispersed repeated sequences including transposons (Delprat *et al*. 2009), retrotransposons (Kupiec and Petes 1988), interspersed repeat sequences (Montgomery *et al*. 1991), or interspersed duplications (Cáceres *et al*. 2007). For example, NAHR between pairs of homologous TEs present in opposite orientations at different positions on a chromosome can lead to inversions of the DNA segment between the two TEs (Delprat *et al*., 2009). Recently, CRISPR has also been used to induce inversions in mammals (Guo *et al*. 2015) and maize (Schwartz *et al*. 2020).

Here we show that DNA transposons, in addition to serving as passive substrates for ectopic recombination, also directly induce inversions via Alternative Transposition reactions. Our results are consistent with a model of RET-induced inversion, in which the ends of two nearby DNA transposons are involved in a single transposition reaction. In this model, two TE copies present in direct orientation will have their adjacent termini in a reversed orientation (i.e., the 5’ end of one TE faces 3’ end of a second TE). Recognition of the terminal sequences of the two TEs by the transposase will lead to an RET event in which the TE termini facing each other attempt to transpose to a genomic target site. Because each TE remains linked to the donor sequences by one un-transposed end, RET results in inversion of a flanking segment, and loss of the fragment originally between the two TEs (Figure 2). Specifically, the DNA segment from one TE end to the new insertion site is inverted. The resulting inversion has TEs present at each breakpoint; one within the inversion and another just outside the second endpoint (Figure 6D). The TE insertion is accompanied by a Target Site Duplications (TSD) flanking the TE termini at the inversion breakpoints. As in standard transposition, the TSD is a result of the staggered cut made by transposase followed by gap-filling and DNA ligation (Lazarow *et al*., 2013).

There are several important differences between inversions resulting from ectopic recombination (NAHR) between two inversely oriented TEs and those caused by RET. First, inversions formed by NAHR will not have a newly generated TSD; instead, the TSDs flanking the internal TE termini will also be inverted, resulting in TEs with (usually) non-identical TSDs. Second, NAHR between two inversely oriented TEs can only flip the intervening segment; whereas, RET can induce inversions of varying lengths on either side of each TE. Third, RET will only operate on Class II TEs that transpose via “cut-and-paste” mechanism, and will not occur with Class I elements that utilize a retro-transposition mechanism. Fourth, RET requires the expression of a DNA transposase and transposition-competent TE termini in appropriate orientation; whereas, NAHR proceeds via the action of host recombination machinery on substrate sequences of sufficient homology and orientation.

The maize *Ac/Ds* system is not the only known system that can cause inversions and other rearrangements. Like *Ac/Ds* elements in maize, the *P-elements* in Drosophila are also known to cause inversions and other chromosomal rearrangements through Alternative Transpositions (Gray *et al*. 1996; Tanaka *et al*. 1997). Other examples of such rearrangements via non-standard transposition include *impala* elements in the fungus Fusarium (Hua-Van *et al*. 2002) and *Sleeping Beauty* transposons in transgenes of mice (Geurts *et al*. 2006).

In addition to RET, the *Ac/Ds* elements can also undergo Sister Chromatid Transposition (SCT) (Zhang, J. and Peterson 2005; Zhang *et al*. 2013). While RET targets TEs on the same chromosome, SCT involves TEs on sister chromatids. After DNA replication, a pair of *Ac* 5’ and 3’ termini in direct orientation can move to an un-replicated region where they can undergo a second round of replication. This results in inverted duplications and Composite Insertions (Wang *et al*. 2020). Both SCT and RET can lead to major rearrangements in the genome. Transposition in the *Ac/Ds* system is non-random (Vollbrecht *et al*. 2010) as *Ac* transposes preferentially into hypomethylated DNA (Kolkman *et al*. 2005) often associated with genic regions (Cowperthwaite *et al*. 2002). This insertion preference likely increases the potential genetic impact of *Ac/Ds-*induced Alternative Transposition events.

### Frequency of Inversions and Other Rearrangements

In a previous study, Yu *et al*. (2011) screened alleles with reverse-oriented *Ac/fAc* insertions in an active *p1* gene for RET-induced loss of function mutants. Out of 100 mutants obtained, 89 were identified to have undergone major structural changes. Approximately half (47 out of 89) were inversions, and the rest were primarily deletions plus some other rearrangements. This result is consistent with the RET model which predicts that inversions and deletions are equally likely to occur, because the outcome is determined by which transposon end (*Ac* or *fAc*) is ligated to which side (*a* or *b*) of the transposition target site. Here, we screened ears from roughly 4000 plants of *p1-wwB54/p1-ww*(*J*) genotype for red kernels indicating putative rearrangements. About 400 unique red kernel events were found and propagated. The red pericarp phenotype was inherited in 97 cases; 83 of these were characterized as rearrangements due to RET. Among these 83, only 14% (12 out of 83) were inversions, 35% (29) were deletions, and 51% (42) were Composite Insertions. The markedly different proportion of inversions recovered here (14%) compared to Yu *et al*. 2011 (53%) is most likely due to the different screens used to detect RET events. The 2011 study began with a functional *p1* gene and selected for loss-of-function events, yielding mostly deletions and inversions; most Composite Insertions would not be detected because they leave the original donor locus intact (Zhang *et al*. 2014; Su *et al*. 2018, 2020). Whereas, this study began with a non-functional *p1* allele, and required gain-of-function (red pericarp sectors). This selection favored recovery of *p2-*expressing alleles caused by inversions and Composite Insertions near *p2* (Su *et al*., 2020). Indeed, all six of the cases described here have inversion breakpoints within 3.5 kb upstream of the *p2* gene. This brings the *p2* promoter to within 10 kb of the *p1* pericarp enhancer (Sidorenko *et al*., 2000), thus activating the *p2* gene in a tissue in which it is not normally expressed.

The six inversion cases described here have no other detectable rearrangements. However, we also obtained seven other cases of inversions which contain other more complicated structural rearrangements. These cases of complex inversions are currently being characterized and will be described elsewhere.

### Effects of Inversions on Fitness

Inversions can have a variety of effects, such as causing position effect variegation of *white* gene in Drosophila (Muller 1930; Levis *et al*. 1985; Lerach *et al*. 2006; Bao *et al*. 2007), suppressing recombination (Jiang *et al*. 2007), and playing a vital role in the evolution of sex chromosomes (Wright *et al*. 2016). Inversions are also associated with local adaptation and reproductive isolation (Lowry and Willis 2010), as many closely related species are thought to have diverged via inversion polymorphisms (Oneal *et al*. 2014; Twyford and Friedman 2015). Inversion of boundary elements may also change higher-order organization in mammalian genomes, due to the directional nature of CTCF binding sites (Guo *et al*. 2015). By altering topologically associated domains (TAD) boundaries, inversions can cause misexpression and disease by changing the relative position of enhancers and their target promoters (Lupiáñez *et al*. 2015; Bompadre and Andrey 2019).

Some inversions can result in major adaptive advantages; for example, the paracentric inversion in *Arabidopsis thaliana* induced by *Vandal* transposon activity is strongly associated with fecundity under drought conditions (Fransz *et al*. 2016). Inversions can even affect the spread of disease: a chromosome 2La inversion in *Anopheles gambiae* is associated with susceptibility of the vector to malaria infection (Riehle *et al*. 2017). Inversions are also involved in local adaptation in teosinte populations (Pyhäjärvi *et al*. 2013). A large (13 Mb) inversion called *Inv4m* found in Mexican highland maize populations affects expression of a large number of genes regulating various developmental and physiological processes contributing to local adaptation to highland environments (Crow *et al*. 2020).

The phlobaphene pigments controlled by the maize *p1* gene are non-essential, and many modern corn varieties lack significant kernel pericarp color. However, a recent study reported that high phlobaphene levels were associated with increased kernel pericarp thickness and reduced mycotoxin contamination when compared to isogenic colorless pericarp lines lacking an active *p1* gene (Landoni *et al*. 2020). Because the *p1* and *p2-*encoded proteins are highly similar and regulate the same flavonoid biosynthetic pathway (Zhang P. *et al*. 2000), similar effects are likely induced by the expression of *p2* in the pericarp. Thus, the transposon-induced inversions identified here may provide an adaptive benefit. Small (< 1 Mb) inversions are difficult to detect by genetic and cytological methods, and so their frequency in plant populations is often unknown. Our results show that even small, cytologically undetectable inversions between linked genes may positively affect fitness. In summary, these findings suggest that Alternative Transposition events may play a critical role in altering gene expression and generating adaptive variation during genome evolution.

## Acknowledgments

We thank Jeremy Schuster and Matthew Johnston for field assistance, and Terry Olson for technical assistance. This research is supported by the USDA National Institute of Food and Agriculture Hatch project number IOW05282, and by State of Iowa funds.

## Supplemental Information

**Figure S1:**
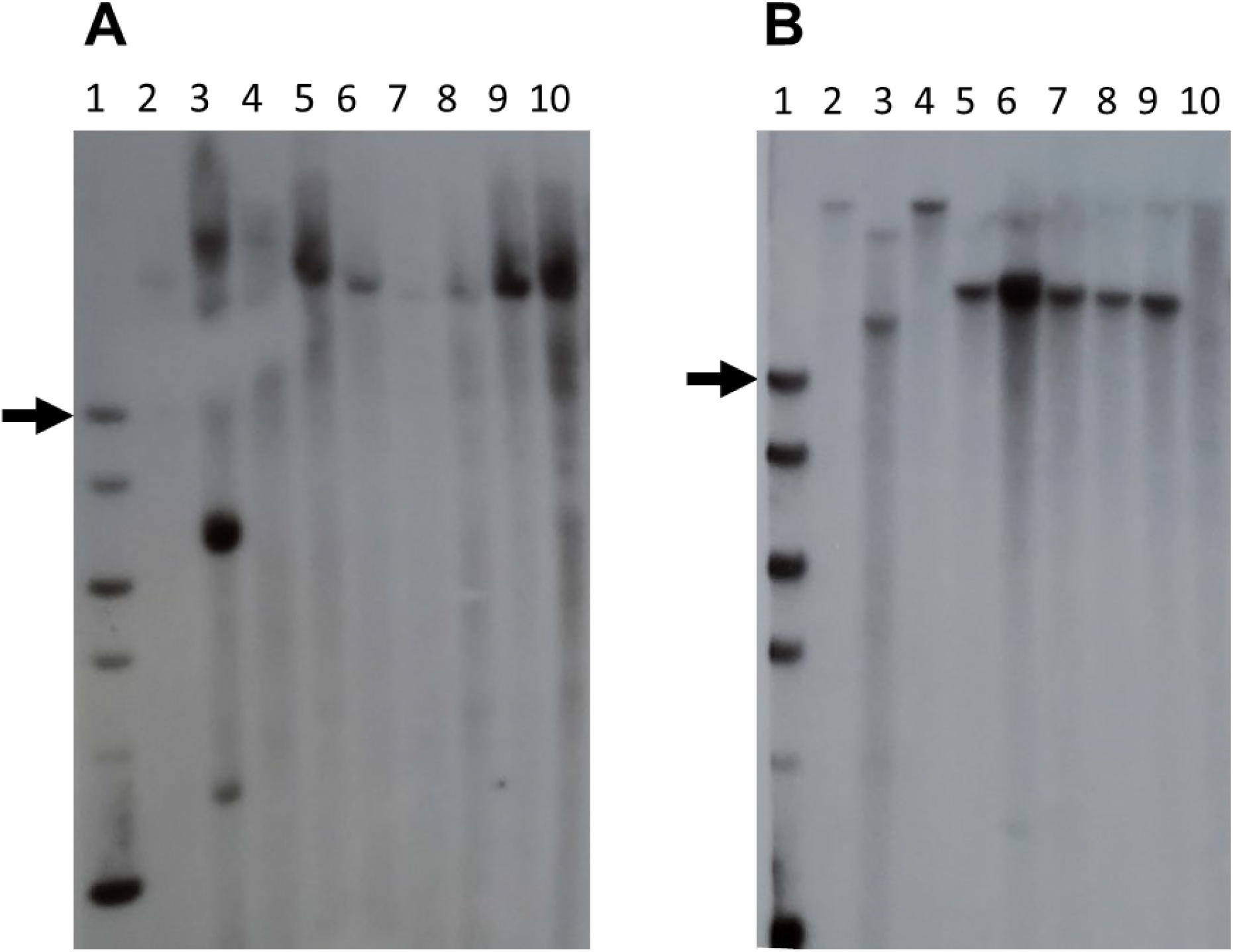
Southern Blot gel images using A) *Hpa*I and B) *Eco*RV restriction enzymes with fragment-15 (within the *p1* enhancer) as a probe. Lane 1, DNA ladder, black arrow points to 10 kb fragment on each gel; Lane 2, J (*p1-ww*); Lane 3, *P1-rr4B2*; Lane 4, *p1-wwB54*; Lane 5, 140; Lane 6, E1; Lane 7, TZ3-4; Lane 8, SP1-18; Lane 9, S25; and Lane 10, TZ2-7.

**Figure S2:**
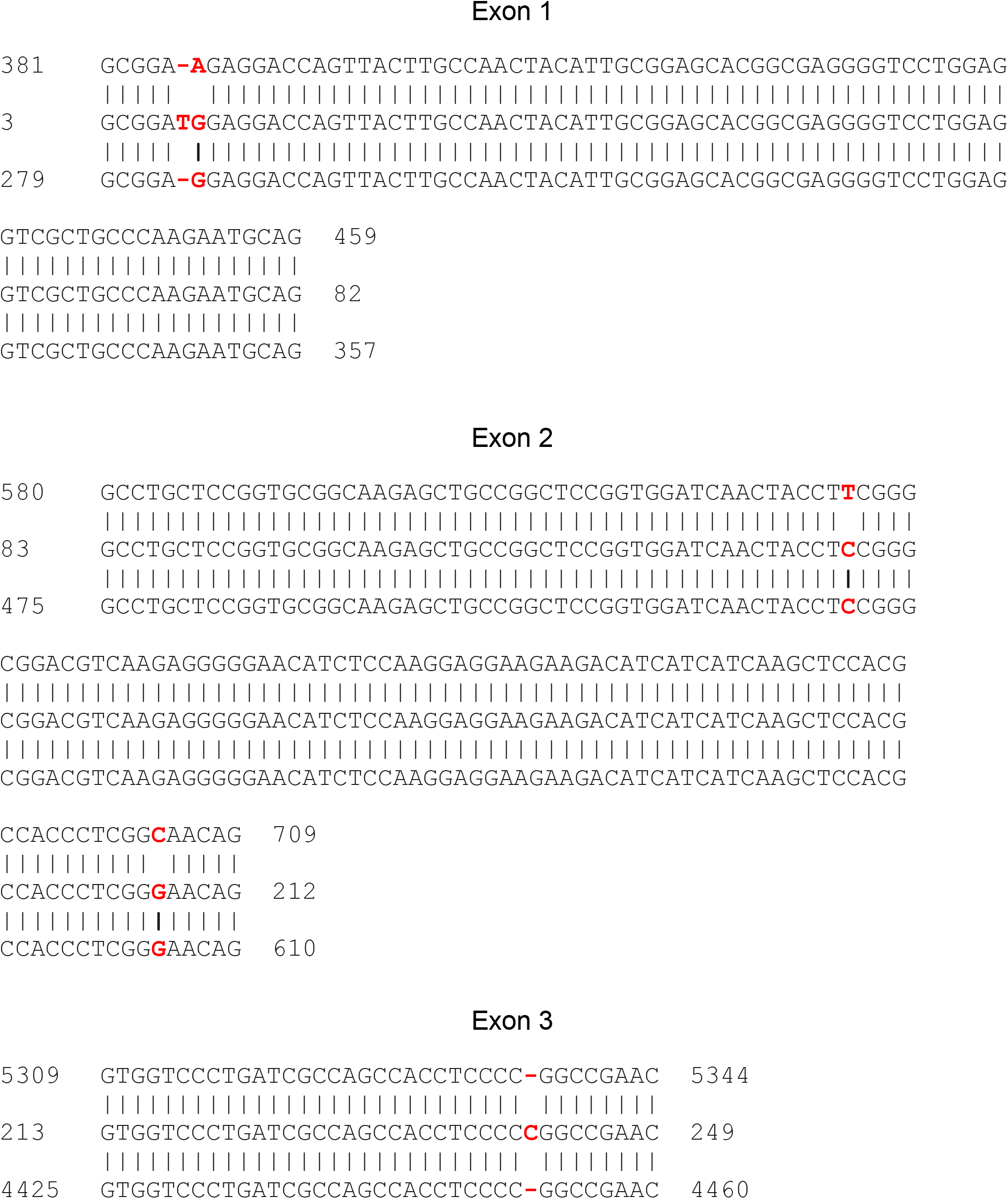
RT-PCR sequence aligned to *p1* and *p2* exons 1, 2 and 3. The middle sequence is RT-PCR product from E1 (one of the inversions), the upper sequence is from *p1* and the lower sequence is from *p2*. At three sites, SNPs in the RT-PCR product match *p2* (lower) but not *p1* (upper). Two additional SNPs in the RT-PCR product likely represent amplification or sequencing artefacts.

**Table S1:**
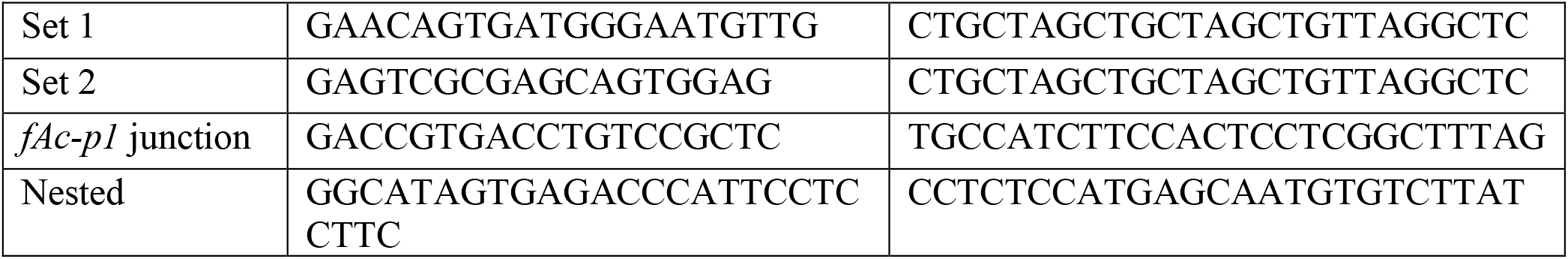
Primers used for screening for inversions.

**Table S2:**
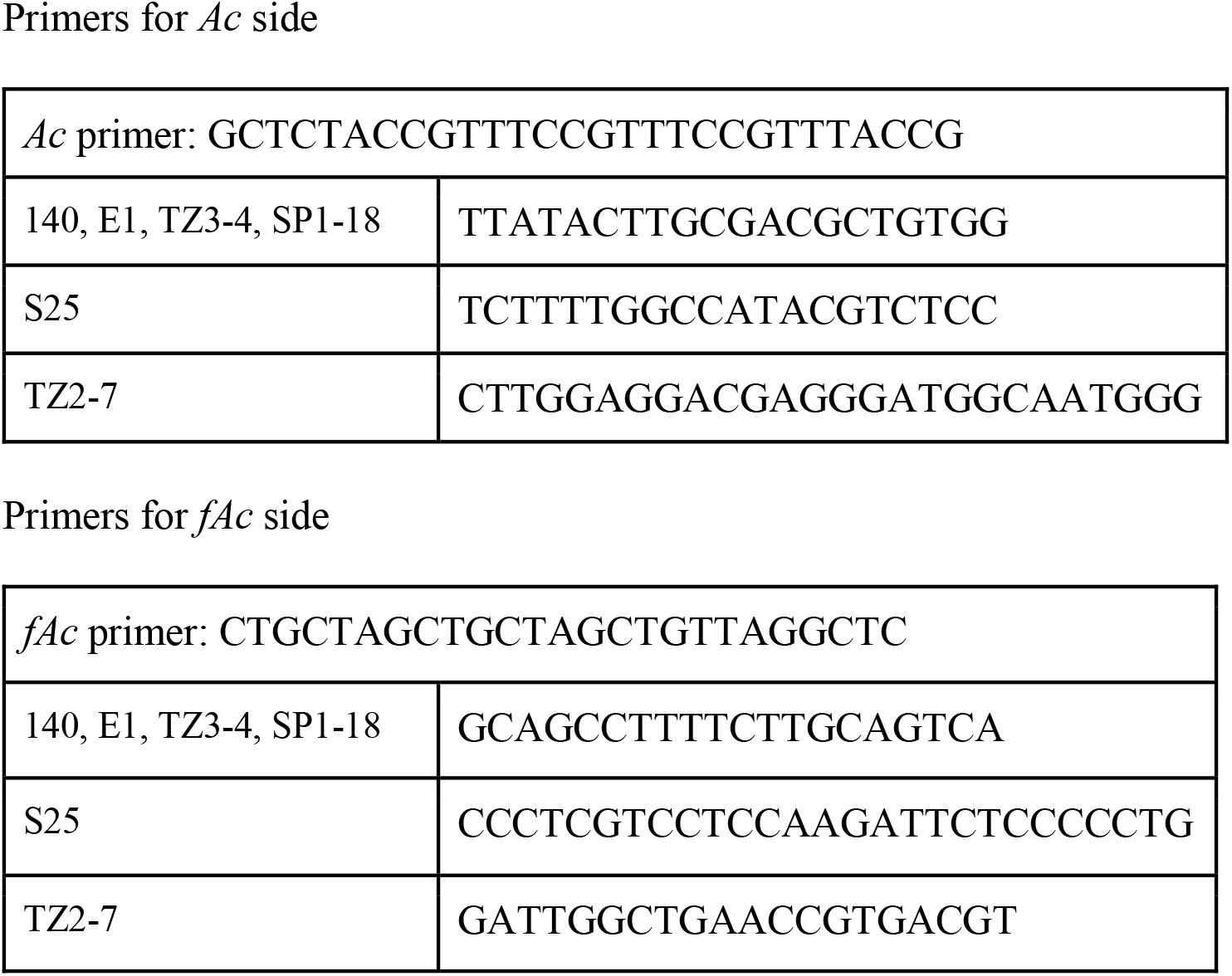
Primers for sequencing inversion endpoints.

**Table S3:**
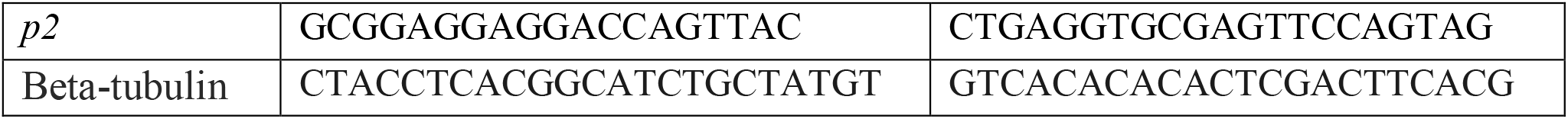
Primers used for RT-PCR.

**Table S4:** Inversion alleles, Target Site Duplications, and relevant distances (in basepairs)

| **Name** | **Target Site Duplication** | **Distance from TSS** | **E-P distance** | **Size of Inverted fragment** |
| --- | --- | --- | --- | --- |
| 140 | CCGGTGGC | 3505 | 8235 | 81800 |
| E1 | CAGCCAGG | 3387 | 8117 | 81682 |
| TZ3-4 | TGTGTAGT | 3376 | 8106 | 81671 |
| SP1-18 | GTCGGGGC | 3203 | 7933 | 81498 |
| S25 | CTCGTCGA | 3070 | 7800 | 81365 |
| TZ2-7 | ATCTCTTC | 2692 | 7422 | 80987 |

